# Evolution of quantitative trait locus hotspots in yeast species

**DOI:** 10.1101/2021.04.07.438839

**Authors:** Emilien Peltier, Sabrina Bibi-Triki, Fabien Dutreux, Claudia Caradec, Anne Friedrich, Bertrand Llorente, Joseph Schacherer

## Abstract

Dissecting the genetic basis of complex trait remains a real challenge. The budding yeast *Saccharomyces cerevisiae* has become a model organism for studying quantitative traits, successfully increasing our knowledge in many aspects. However, the exploration of the genotype-phenotype relationship in non-model yeast species could provide a deeper insight into the genetic basis of complex traits. Here, we have studied this relationship in the *Lachancea waltii* species which diverged from the *S. cerevisiae* lineage prior to the whole-genome duplication. By performing linkage mapping analyses in this species, we identified 86 quantitative trait loci (QTL) affecting growth fitness in a large number of conditions. The distribution of these loci across the genome has revealed two major QTL hotspots. A first hotspot corresponds to a general fitness QTL, impacting a wide range of conditions. By contrast, the second hotspot highlighted a fitness trade-off with a disadvantageous allele for drug-free conditions which proved to be advantageous in the presence of several drugs. Finally, the comparison of the detected QTL in *L. waltii* with those which had been previously identified for the same traits in a closely related species, *Lachancea kluyveri*, clearly revealed the absence of interspecific conservation of these loci. Altogether, our results expand our knowledge on the variation of the QTL landscape across different yeast species.

## Introduction

Understanding how the genetic diversity generates the impressive phenotypic variation observed at the species level is one of the greatest challenges in biology. A better view of the genotype-phenotype relationship also provides helpful information in health research (Minikel et al. 2020), the food industry (McCouch 2004; Marullo et al. 2006; Sharmaa et al. 2015), as well as on evolution and adaptation mechanisms (Olson-Manning, Wagner, and Mitchell-Olds 2012). Exploring the genetic basis of quantitative traits is not a trivial task as it may involve a large number of alleles with different effect size and interacting with each other as well as with the environment (Lynch and Walsh 1998). Two main strategies make it possible to link phenotypic variation to specific loci: genome-wide association studies (GWAS) which use large samples of individuals, and linkage analyses that increase statistical power by generating recombinant offspring (Lynch and Walsh 1998).

The *Saccharomyces cerevisiae* yeast appeared very early to be an excellent model to perform linkage analyses (Steinmetz et al. 2002; Brem et al. 2002) and a large number of QTL mapping studies have been conducted to better understand epistasis (Sinha et al. 2006), missing heritability (Bloom et al. 2013), gene-environment interactions (Smith and Kruglyak 2008; Bhatia et al. 2014; Yadav, Dhole, and Sinha 2016; Peltier et al. 2018), the impact of rare variants (Fournier et al. 2019; Bloom et al. 2019) and to the accumulation of hundreds of SNPs of technological interest (Peltier et al. 2019). However, comprehensive understanding of phenotypic diversity would benefit from studying the genotype-phenotype relationship in various species. Only a few QTL mapping studies have been performed on other yeast species (Clément‐Ziza et al. 2014; Sigwalt et al. 2016; Brion et al. 2020). Saccharomycotina yeasts span a broad evolutionary scale shaped by different forces as they have different environmental niches, life cycle, mating type system, ploidy level and not all have been subjected to artificial selection through domestication such as *S. cerevisiae* (Dujon 2006; Peter and Schacherer 2016). Exploring phenotype-genotype relationship in additional yeast species is therefore of great interest.

In this context, we explored the phenotype-genotype relationship in another yeast species, *Lachancea waltii*. This yeast has a ∼11 Mb genome distributed in eight chromosomes and diverged from the *S. cerevisiae* lineage prior to the whole-genome duplication event, estimated to have occurred more than 100 Ma (Kellis, Birren, and Lander 2004). The ecological niche of *L. waltii* is poorly documented and some strains have been isolated from tree exudates, insects or lake water. Unlike other species such as *Lachancea thermotolerans, L. waltii* has never been found associated with human activities (Porter, Divol, and Setati 2019). In the laboratory, *L. waltii* grows in standard *S. cerevisiae* rich media and is sensitive to drugs commonly used for *S. cerevisiae* (Di Rienzi et al. 2011). Mating type and silent cassettes are present on the left arm of chromosome C while neither *HO* gene nor *HO* recognition sequences are found (Fabre et al. 2005). Although rare *HO*-independent mating type switches are suspected, *L. waltii* appears to be a stable, vegetatively growing haploid (Di Rienzi et al. 2011). Mating is suspected to be rare on rich media due to the absence of pheromone response and crosses are successfully generated using selective markers leading to stable diploids (Di Rienzi et al. 2011). These life cycle properties therefore offer the opportunity to perform quantitative genetic studies on this species with a classical QTL mapping design by generating a large recombinant offspring from a parental cross.

To dissect the genetic basis for variation in fitness under a broad set of environmental conditions in *L. waltii*, we performed linkage mapping on a large population of segregants. To this end, two natural isolates were crossed to generate a set of 421 segregants, which were phenotyped for their ability to grow under a total of 48 environmental conditions. The phenotypic variation within this population is highly heritable and displays mainly a normal distribution, characteristic of quantitative traits. The genomes of this set of segregants were completely sequenced and the parental segregating alleles were used as markers to perform linkage analysis. We found a total of 86 QTL, mostly grouped into two major hotspots. Their impact on the phenotypic landscape were examined and revealed a general growth as well as a multidrug resistance QTL. Comparison of the detected QTL in *L. waltii* with those we previously determined in a closely related species, *L. kluyveri*, clearly showed the absence of overlap between the QTL landscapes. Unlike the *L. waltii* species, *L. kluyveri* is characterized by a major QTL hotspot showing an association between a large number of traits and the mating-type locus due to the absence of recombination in this specific region. And this situation leads to the presence of a sexual dimorphism which is absent in *L. waltii* and consequently highlighting a divergent evolutionary trajectory in the *Lachancea* genus. Overall, our results provide insight into the evolution of the genetic architecture of quantitative traits across yeast species.

## Results

### Lachancea waltii mapping population and fitness landscape

To identify the genetic basis underlying the phenotypic diversity in *L. waltii* species, we selected two isolates, LA128 and LA136, as parental strains (Table S1). While LA128 was isolated from the gall of a red oak tree, LA136 was isolated from the black knot of *Prunus virginiana* but both strains were isolated in Canada. After whole genome sequencing, a total of 64,628 confident SNPs discriminating LA128 and LA136 were identified, leading to a genetic divergence of 0.59 %. The LA128 and LA136 stable haploid strains were mated to generate a LA128/LA136 hybrid (Figure S1). This diploid was sporulated and a total of 421 F1 offspring were isolated. At each step, the ploidy level was checked by flow cytometry. These segregants were genotyped by whole-genome sequencing attributing for each individual their SNP parental inheritance. A subset of 5,542 SNPs assigned with confidence in over 94% of the progeny and distributed homogeneously along the genome were selected for use as genetic markers for linkage analysis (Table S2). The correct 2: 2 segregation of these markers shows that this mapping population is suitable for performing QTL mapping

Then, we sought to capture the phenotypic diversity of this population and fitness variation was assessed by measuring colony growth on different media (Table S3). The impact of 48 conditions on growth fitness was tested, including 16 drugs / compounds with multiple concentration levels, two non-glucose carbon sources (galactose and glycerol) and two temperatures (14°C *vs*. 24°C) (Table S4). Briefly, the compounds tested are antifungal compounds, ions, metals, solvents or toxic compounds described in the literature to have an impact on yeast fitness. We first studied the phenotypic distribution and variance of the offspring. While most traits display a complex distribution, some specific media highlight bimodal distributions in at least one concentration level (benomyl 50 -100 µg/mL, fluconazole 1 µg/mL, glucose 10 %, CuSO4 0.1 mM, formamide 1.5 %, 6-azauracil 100 µg/mL, bafilomycin 0.375 µg/mL and caffeine 10 mM). Representative examples of phenotypic distribution are shown in Figure 1A. For all traits, parental strains show significant differences in fitness for at least one concentration level. The most striking differences are observed for 4-nitroquinoline 1-oxide (1 μg/mL) and galactose (2 %) conditions in which almost no growth is observed in LA128 while LA136 is able to grow. Overall, LA128 has a significative higher fitness in 18 conditions compared to 21 conditions for LA136, showing that the genetic basis of fitness is different under the conditions tested and that favorable alleles are distributed in both parents. The rank of the parents can change depending on the level of compound concentration level (see example of fluconazole in Figure 1A) illustrating the presence of genotype-environment interaction (GxE). To have a view of the phenotypic diversity in this offspring, the phenotypic variance by conditions was estimated. As expected, we observed a large phenotypic variance under most conditions. While some conditions show high phenotypic variance (*e*.*g*. glucose 10 %, 6-azauracil 100 µg/mL and caffeine 10 mM), others exhibit very low variance, mainly because the drug concentration level was too high and allowed only low growth (*e*.*g*. sodium dodecyl sulfate 0.025 % -0.05 %, and 4-nitroquinoline 1-oxide 3 µg/mL). The broad-sense heritability (*H*^*2*^) was computed to assess the fraction of the phenotypic variance caused by genetic factors (Figure 1B-C). The heritability is 0.69 on average, with 80 % of the traits having a heritability greater than 0.5. Taken together, these results indicate that the phenotypic variance observed in the offspring is primarily caused by heritable genetic factors and can be mapped with our experimental design.

**Figure 1.**
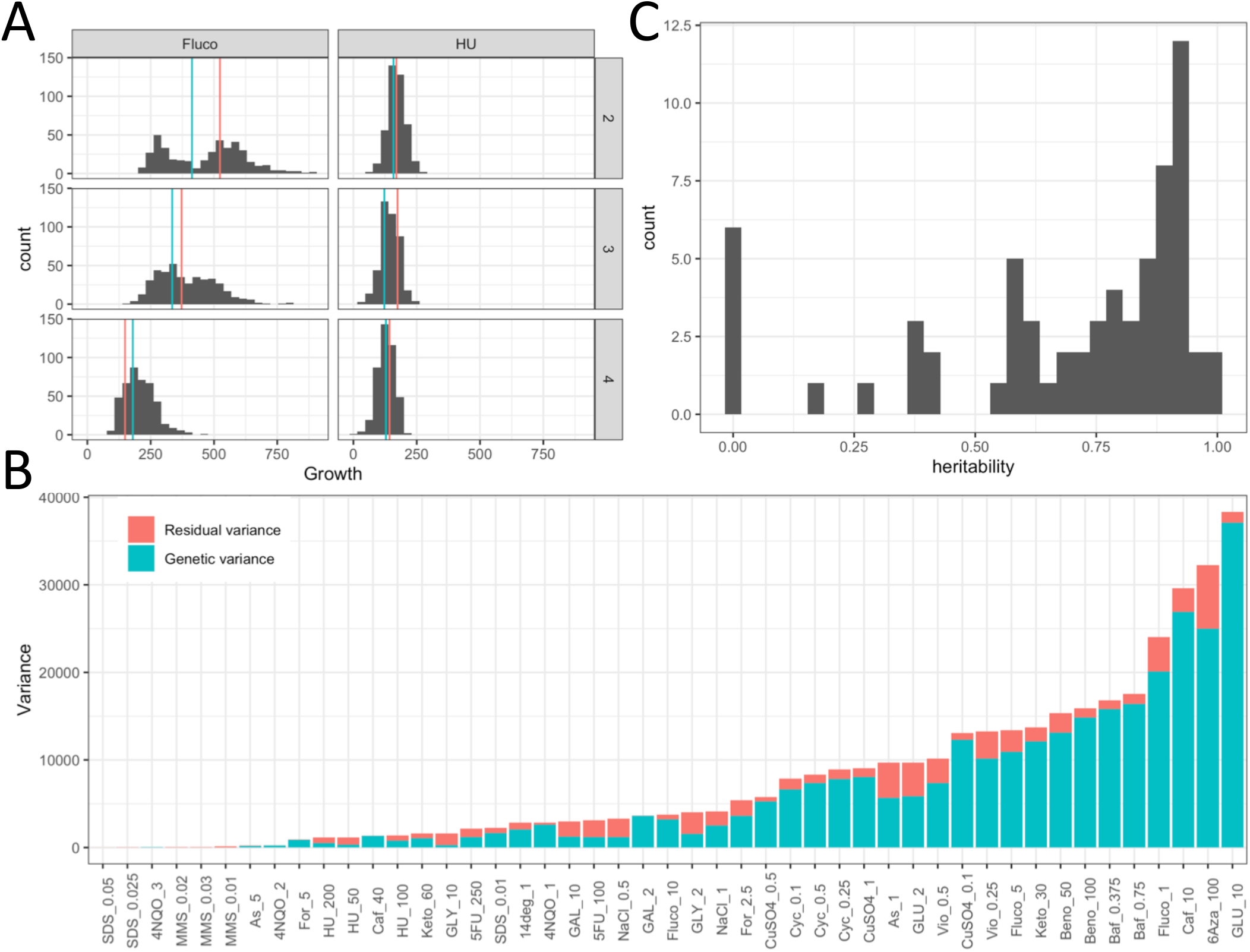
Distribution, genetic variance and heritability of the traits. **(A)** Distribution of growth fitness for F1 offspring. Red and blue horizontal lines represent the parental value of LA128 and LA136, respectively. Panel 2, 3 and 4 represent an increase in the concentration level with 1 -5 -10 μg/mL of fluconazole and 20 -100 -200 µg/mL for hydroxyurea. **(B)** Variance in the F1 offspring per media. The total variance is composed of the genetic variance and residual variance. **(C)** Distribution of the *h*^*2*^ estimated for the 48 media.

### Two major QTL hotspots shape the *L. waltii* fitness landscape

The genotypic and phenotypic data obtained for the offspring were used to perform linkage mapping analysis. Here, a trait was considered to be the colony growth in a medium with a specific drug or carbon source and we consequently studied a total of 22 traits. For each trait, the concentrations of compounds or the different temperature levels were considered as environmental variables. We then identified QTL and the variation of their effect according to the concentration have been evaluated (GxE). This linkage analysis led to the identification of 86 QTL with a false discovery rate of 1 % (Figure 2A and Table S5). An average of 4 QTL per trait was found with at least one QTL per trait and a maximum of 7 QTL identified for fitness in the presence of benomyl, cycloheximide and methyl viologen. We then looked at the distribution of the QTL across the genome. Since many QTL significantly span multiple markers and several kb, their position has been assigned to the marker with the highest significance. Using a 50 kb window, the number of QTL per window was determined. This analysis revealed a total of 86 QTL distributed across 16 different loci, with 6 of them impacting at least 6 different traits (Figure 2B). Strikingly, two of these pleiotropic loci are QTL hotspots that affect almost every trait. These two hotspots are mapped in the chromosome E, at positions 87402 and 810257 (hereafter named E_87 and E_810) and have an effect on 19 and 20 traits, respectively. To assess the extent to which phenotypic diversity is explained by the identified QTL, the variance explained by each QTL was determined with analysis of variance (ANOVA). Each QTL explains 8 % of the variance on average. The distribution of the variance explained displays a typical L-shape distribution often found in QTL mapping analysis, with most of the QTL explaining low percentage of variation and few major QTL explaining high part of the variation (Figure S2). Indeed, 85 % of the QTL explain less than 15 % of the variance. The major QTL explaining a significant proportion of the variance are the two hotspots E_87 and E_810 mapped in almost all of the traits (19 and 20, respectively) (Figure 2B). To assess the phenotypic impact of these two major QTL, a principal component analysis with the phenotypic dataset was performed (Figure 3A). The two first dimensions account for 48 % of the variance and clearly discriminate 4 groups according to E_87 and E_810 allele inheritance. The segregants that inherited the parental genotypes are located in two groups with their corresponding parent while the other two groups which inherited recombined genotypes are clearly segregating separately. This is explained by significant phenotypic transgression for most traits, with the lowest fitness segregants being those that inherited the deleterious E_87^LA136^ and E_810^LA128^ combination and the highest fitness segregants being those that inherited the favorable E_87^LA128^ and E_810^LA136^ combination. A representative example of the transgression found is shown in the copper sulfate condition (0.5 mM) (Figure 3B). Another ANOVA was performed to identify potential interactions between these two major QTL and no significant interaction was found, showing strong additivity between E_810 and E_87 (Figure 3C). The high individual effect of these major QTL and their additivity explain their drastic impact on the phenotypic landscape. Moreover, the fact that parental strains have alleles of opposite effect explains the level of transgression observed in the offspring. Overall, this analysis showed that the phenotypic diversity of this cross is mainly shaped by two major additive QTL whose effect is revealed by meiosis leading to transgressive individuals.

**Figure 2.**
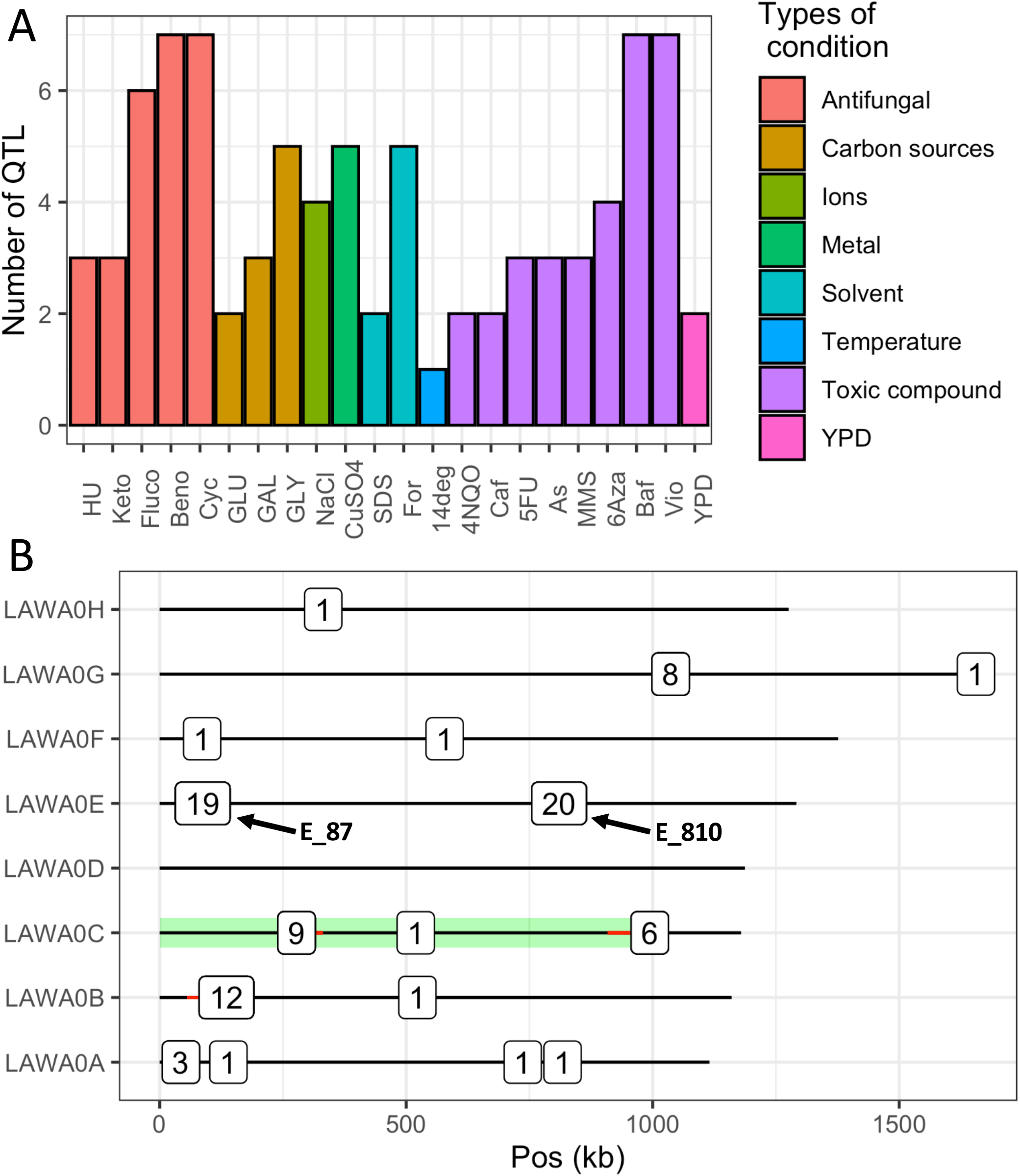
Overview of the identified QTL. (**A**) Number of QTL identified per trait. **(B**) Genomic location of the QTL identified for all traits. Each square indicates at least one QTL, the number in each square indicate the number of QTL identified in the same 50 kb window. When present, the red line indicates window limit. Green shadow represents the left arm of chromosome C.

**Figure 3.**
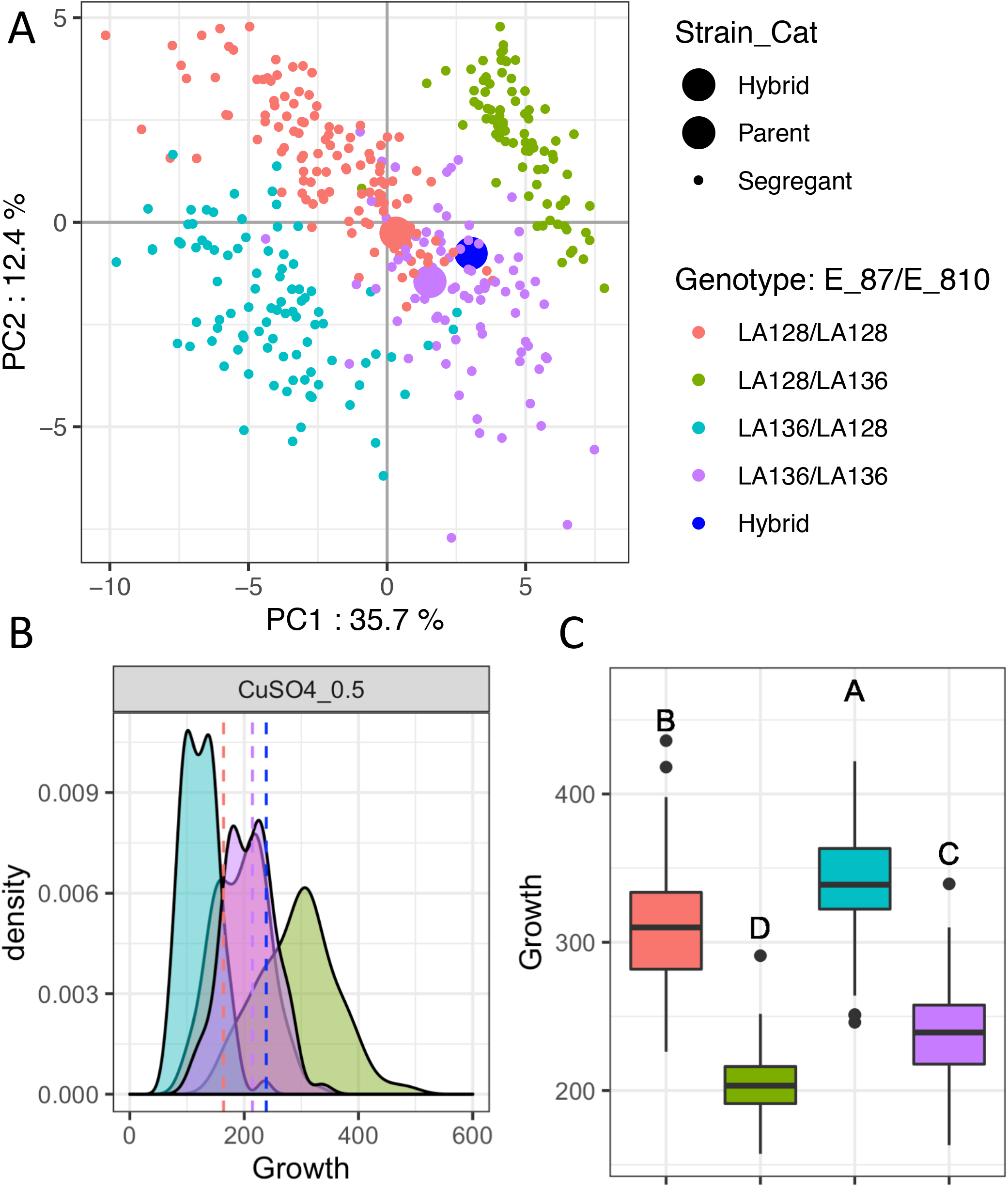
Overall effect of the two QTL hotspots detected. **(A)** PCA of all the fitness measure for the F1 population, parental strains and hybrid. The color by genotype for E_87 and E_810 QTL is the same for all panels. **(B)** Distribution of the fitness of the segregants in copper sulfate 0.5 mM according to their genotype. Vertical dashed lines represent the parental strains and the hybrid. **(C**) Additive effect of E_87 and E_810. The boxplot represents the segregant fitness in all media according to their genotype. A different letter indicates a significant difference (Tukey Honest Significant Differences, confidence level = 0.95).

### Identification of a general growth QTL with high level of interaction according to media complexity

We next studied in more detail the effect of the two main QTL, *i*.*e*. E_810 and E_87. The E_810 hotspot reaches the significance threshold in 20 out of the 22 traits. It explains 27 % of the variation on average, with a maximum of 76 % in glucose 10 % (Figure 4A). The effect is exacerbated in less stringent media with almost no effect (6 % of variance explained on average) in the 18 more stringent media (Figure 4B-C). Therefore, its effect is amplified in any condition without drugs (*i*.*e*. 2% and 10% glucose) or with a low level of concentration. For these conditions, E_810 explains perfectly the bimodal distributions identified. Interestingly, this QTL has only a small effect on the non-glucose carbon sources such as glycerol and galactose (Figure 4A). While the low effect in galactose can be attributed to the general low fitness for this carbon source, this is not the case for glycerol. This shows an important GxE interaction between E_810 and the carbon source, the E_810 impact being greater with glucose. The interaction is more complex when compared with YPD in which E_810 clearly has no significant effect (Figure 4A). The environmental factor which generates this striking GxE effect cannot be clearly determined because the synthetic complete and YPD media composition differs in many aspects. The variation in the expression of phenotypes according to complex yeast media and synthetic complete media has already been reported in the literature (Janitor and Šubík 1993; Supek et al. 1996; Kucejova et al. 2005; Abelovska et al. 2007). Therefore, this important GxE interaction can be generated by a substantial difference in the concentration of key elements involved in the E_810 effect. We concluded that E_810 hotspot is a general growth QTL whose drastic impact is revealed by minimum media.

**Figure 4.**
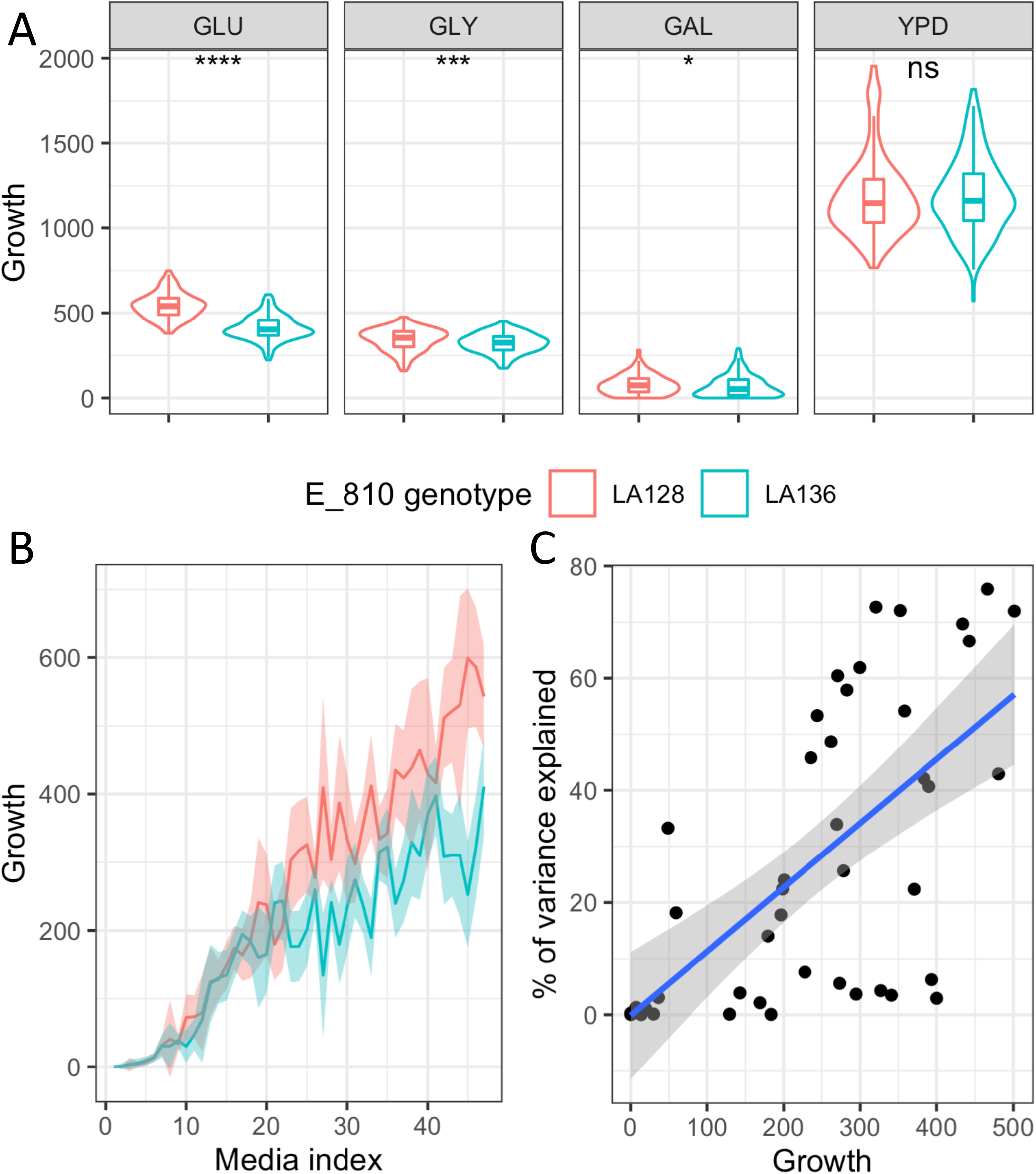
Effect of the E_810 QTL hotspot. **(A**) Grotwth of F1 offspring according to E_810 inheritance and carbon source glucose 2 % (GLU), glycerol 2 % (GLY), galactose 2 % (GAL) or media base Sc vs YPD. Wilcoxon–Mann–Whitney test was applied to assess the significance of the phenotypic difference between the relevant pairs. The level of significance is indicated as follows: ns: not significant, * *p* ≤ 0.1, ** *p* ≤ 0.05, *** *p* ≤ 0.01. Linkage analysis, which is more stringent than Wilcoxon–Mann–Whitney test due to multiple tests correction by permutations doesn’t link significantly E_810 to galactose. **(B**) Reaction norm of F1 offspring according to E_810 inheritance in all 48 media. Media are sorted according to the average growth of the F1 offspring. **(C**) Average growth per medium according to the variance explained by E_810. Each dot represents a condition for which E_810 is identified as QTL.

### Identification of a multidrug resistance QTL highlighting an evolutionary trade-off

We then investigated in more detail the effect of E_87 in the F1 offspring. This QTL is detected for 19 out of the 22 traits. It has a lower effect than E_810, explaining 12 % of the variation on average, with a maximum of 45 % in hydroxyurea (100 µg/mL). Interestingly, while the E_87^LA136^ allele confers better fitness under most conditions (n = 43), its effect is deleterious under certain specific conditions (n = 9) (Figure 5A). In fact, we can see that E_87^LA128^ appears as a beneficial allele when the drug concentration increases in media with hydroxyurea, 5-fluorouracile, sodium meta-arsenite and caffeine (Figure 5B). This QTL also represents a trade-off with the resistant allele decreasing fitness under most conditions, but improving fitness under specific conditions conferring drug resistance. The fitness cost of drug resistance variants is frequently described in literature and therefore not surprising (Melnyk, Wong, and Kassen 2015; Maharjan and Ferenci 2017).

**Figure 5.**
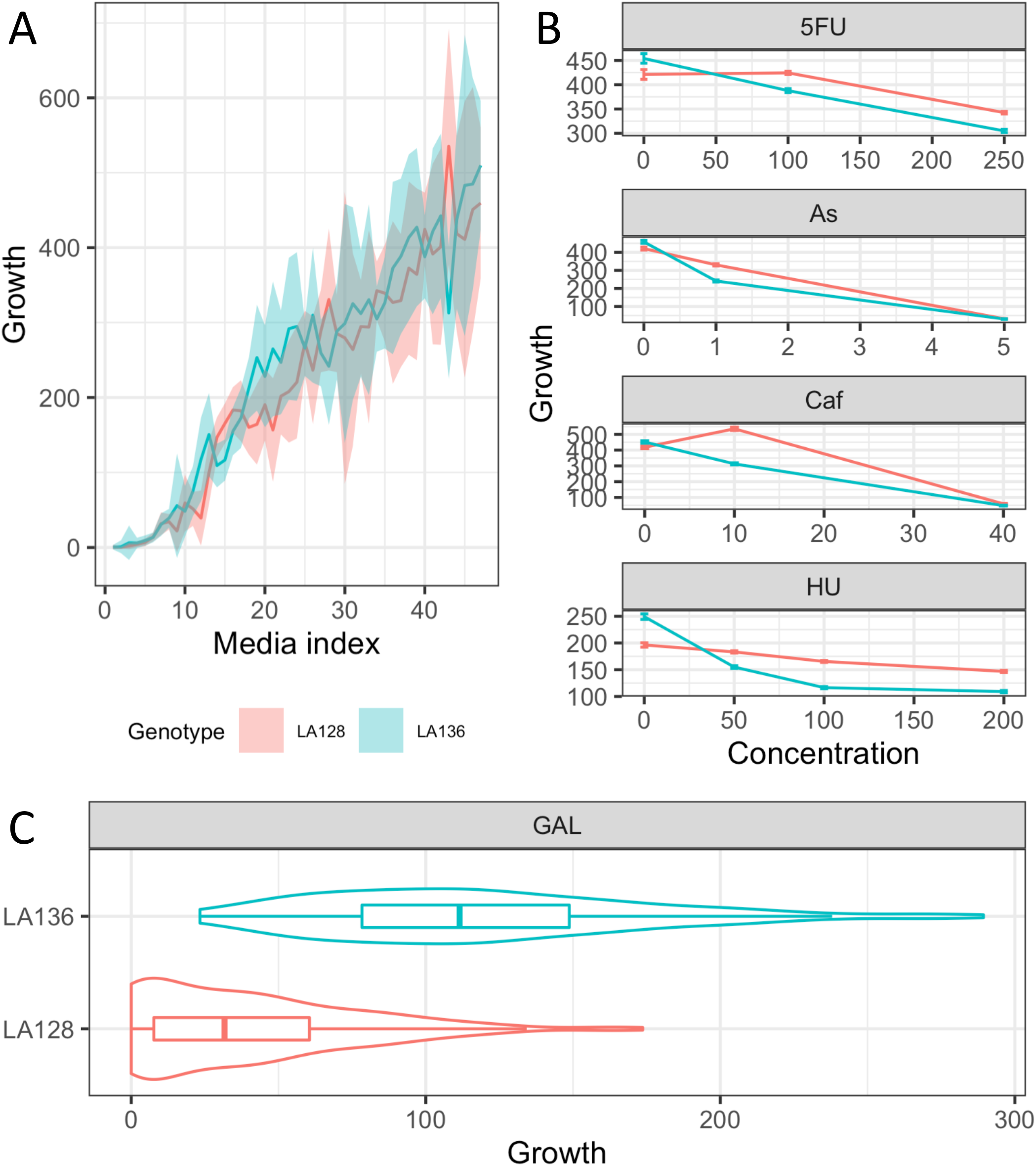
Effect of the E_87 QTL hotspot. **(A)** Reaction norm of F1 offspring according to E_810 inheritance in the 48 media. Line media are sorted according to the average growth of F1 offspring. **(B)** Reaction norm of F1 offspring for some specific traits. **(C)** Growth of F1 offspring according to E_810 inheritance in galactose 2 %.

The targets of the drugs for which E_87 has been detected are multiple, including DNA synthesis and repair, oxidative stress, and signal transduction. The genetic basis underlying this QTL can be linked either to different independent genetic factors each having an impact on a specific drug, or to a pleiotropic genetic factor giving resistance against several drugs. Interestingly, we have identified *LAWA0E00386g* as a good candidate gene in this locus, which is an ortholog of the well characterized *PDR15 S. cerevisiae* gene encoding for a multidrug transporter and general stress response factor implicated in cellular detoxification (Wolfgert, Manmun, and Kuchler 2004). In this gene, 8 non-synonymous SNPs discriminating parental sequences were found and may impact its function. To conclude, we hypothesize that E_87 is related to a multidrug-resistant allelic variant resulting in a fitness cost in drug-free condition.

The effect of the E_87 locus is not limited to drugs-containing media and also has an effect in galactose media (2 and 10 %) (Figure 5C). Under these conditions, this QTL has a strong effect, clearly discriminating growing of non-growing strains (41 % of the variance is explained in galactose 2 %). All of the segregants that inherited the E_87^LA136^ allele are able to grow with galactose as the sole carbon source while almost half of the E_87^LA128^ show no growth. As the E_87 locus has a pleiotropic effect on drug resistance and galactose assimilation, we can assume that different genetic factors may explain these unrelated traits. By look at this QTL, we identified the presence of *LAWA0E00342g* which is annotated as similar to *S. cerevisiae GAL2* gene. In *S. cerevisiae*, this gene encodes for a galactose permease (Boles, De Jong-Gubbels, and Pronk 1998) and can therefore be a good candidate gene to explain this difference regarding the assimilation of galactose. Although no non-synonymous SNPs were identified between parental strains, the differences found in the promoter region could explain this phenotypic variation. Surprisingly, the *L. waltii* species was previously reported as unable to assimilate galactose (Porter, Divol, and Setati 2019). In our study, two other QTL with a smaller impact (2-4 % of the variance explained) were also identified for their effect on galactose. This illustrates the presence of natural allelic variants in the *L. waltii* species allowing the assimilation of galactose.

### Absence of conservation of the QTL hotspots across closely related species

To explore whether or not the QTL landscape is conserved across yeast species, we compared the QTL detected here in *L. waltii* with the one of a related species, *L. kluyveri*. We took advantage of our previous work which identified QTL with the same experimental design and similar conditions in *L. kluyveri* (Brion et al. 2020). In this study, 196 F1 segregants were used to perform linkage analysis under 64 conditions, 25 of them being similar or identical to the traits measured in the present work on *L. waltii* (conditions using galactose, glycerol, glucose, sodium chloride, copper sulfate, sodium dodecyl sulfate, formamide, sodium meta-arsenite, methyl viologen, caffeine, 6-azauracil, hydroxyurea, fluconazole, benomyl and cycloheximide). These two datasets therefore provide an excellent opportunity to compare the QTL landscape between two closely related species. With the exception of the 560 first kb of the left arm of the chromosome C (C-left), the synteny between *L. waltii* and *L. kluyveri* is poorly conserved (Vakirlis et al. 2016). To properly compare the QTL landscape and see any possible common QTL, genomic alignment was performed and 203 syntenic blocks of 55 kb on average were identified (Figure 6). In total five syntenic blocks containing at least one QTL in each species were identified. This number is expected by chance under random distribution, showing the absence of conservation of the overall QTL landscape (Pearson’s Chi-squared Test, pval > 0.1).

**Figure 6.** QTL landscape comparison. The genomes of *L. waltii* (LAWA) and *L. kluyveri* (LAKLU) are shown. The vertical red lines represent the boundaries of the chromosomes and letters represents the names of the chromosome. The number of QTL identified per one kb window is indicated and identified by a triangle at the best marker position. All shared synteny blocks that contain at least one best QTL marker in each species are represented by the purple lines and linked to their corresponding position in the other genome by dash lines.

We further investigated the QTL located on shared syntenic blocks to identify the potential presence of any interspecies QTL conservation. We examined a first QTL identified in *L. kluyveri* on the chromosome A and controlling growth on benomyl (0.2 g/L). This QTL is located in the same block of synteny as two QTL identified in *L. waltii* controlling fitness in glycerol (2 %) and galactose (2 %). These QTL are located in the same syntenic block but they are separated by at least 80 kb and control different traits. It is therefore unlikely that the genetic factors behind these QTL are the same. In addition, the two E_87 and E_810 QTL hotspots detected in *L. waltii* are also located in shared syntenic blocks located on the D and H chromosomes of the *L. kluyveri* species. Nevertheless, the *L. kluyveri* QTL has an impact on a limited number of traits (one and two, respectively) and therefore it is again unlikely that the same genetic factors are involved. Overall, no interspecific QTL were identified, confirming the lack of conservation of the QTL landscape between these two species.

Finally, a striking feature of the *L. kluyveri* QTL landscape previously highlighted is the linkage of the whole C-left region with a QTL hotspot impacting 23 traits (Brion et al. 2020). The unusual size of this QTL, that covers a region of a 1 Mb, is explained by the absence of meiotic recombination of the *L. kluyveri* C-left, maintaining the whole arm in genetic linkage (Brion et al. 2017). Interestingly the C-left region contains the *MAT* locus and therefore this lack of recombination plus the presence of a QTL hotspot impacting a broad range of traits revealed a unique case of sexual dimorphism in budding yeast. The C-left synteny is well conserved between the *L. kluyveri* and *L. waltii* species, including the *MAT* locus and the two silent loci *HML*/*HMR*. However, the C-left region in *L. kluyveri* also corresponds to a large GC-rich introgressed region and this is not the case for *L. waltii* (Payen et al. 2009). A total of 16 QTL distributed across 3 distinct loci were detected and no QTL hotspot was found on the C-left in the *L. waltii* species (Figure 2B). This comparison shows that the sexual dimorphism present in *L. kluyveri* is not a general characteristic of the whole *Lachancea* genus which rather diverged in different evolutionary strategies.

## Conclusion

This work presents the first linkage analysis carried out in the *L. waltii* species, demonstrating its suitability to experimental design of classic QTL mapping and extending our knowledge on genetic architecture of quantitative traits to a new yeast species. This analysis led to the identification of 86 QTL, most of them being distributed in two major hotspots. Identification of QTL hotspots is quite common in many species (Mozhui et al. 2008; Ambroset et al. 2011; Sukumaran, Reynolds, and Sansaloni 2018; Larson et al. 2016; Rae et al. 2009). Their identification is almost ubiquitous for eQTL where major regulators or transcription factors may impact several transcripts (Breitling et al. 2008). This high number of QTL identified within the same locus is explained by the non-exclusive combination of two parameters: genetic factors with a pleiotropic effect and multiple genetic factors genetically linked. As all the traits measured in this work correspond to colony growth in different media, they are not completely unrelated. Therefore, a pleiotropic genetic factor impacting fitness under a wide range of growing conditions would be mapped for several traits and this is more likely the case for E_810. This is supported by the fact that the advantageous allele of the multiple QTL associated with this hotspot systematically comes from the same parent. Such hotspots have already been described in *S. cerevisiae* for traits involving the production of related compounds or fermentation traits in an oenological context (Eder et al. 2018; Ambroset et al. 2011). In contrast, the second hotspot, E_87, affects fewer related traits by controlling multiple drug resistance and galactose uptake, so the presence of distinct genetic factors in linkage disequilibrium is more likely.

Our study also allowed us to compare the QTL detected in *L. waltii* and *L. kluyveri* species, showing the lack of conservation of the QTL landscape through related species. Certainly, this comparison is not extensive because it is limited to two species with a cross each. However, some important changes in genetic architecture of quantitative traits, such as sexual dimorphism, can be identified and extended to the whole species. Indeed, a hotspot identified in *L. kluyveri* was linked to an important sexual dimorphism, being the mark of a particular evolutionary strategy, never described in budding yeast before (Brion et al. 2020). The lack of conservation of such a hotspot in *L. waltii* showed an adaptive divergence within the *Lachancea* genus. It would obviously be interesting to extent such linkage analyses to a larger number of yeast species in order to have a broader view of the evolution of the architecture of quantitative traits. Indeed, yeast covers a wide evolutionary range shaped by different forces such as environmental niches, life cycle, mating type system, ploidy level which could impact the architecture of quantitative traits.

## Material and Methods

### Yeast strains and media

Yeast strains used in this study are described in Table S1. Strains were grown in standard YPD medium (yeast extract 1% peptone 2% glucose 2%) supplemented with G418 (200 μg/mL) or nourseothricin (100 μg/mL) and agar 20 g/L for solid medium at 30 °C. The *HIS3*::*KanMX* phenotype was verified on SC-His-Trp-Ura medium (yeast nitrogen base with ammonium sulfate 6.7 g/L, SC-His-Trp-Ura amino acid mixture 1.74 g/L, dextrose 20 g/L, and agar 20 g/L). Mating and sporulation were performed on DYM medium (yeast extract 0.3 g/L, malt extract 0.3 g/L, peptones 0.5 g/L, dextrose 1 g/L and agar 20 g/L) at 22°C. Phenotyping experiments were performed on synthetic complete (SC) medium (yeast nitrogen base with ammonium sulfate 6.7 g/L, SC amino acid mixture 2 g/L, sugar 20 g/L, agar 20 g/L) supplemented with specific compounds (Table S4).

### Generation of parental strains

Stable haploid parental strains were obtained by replacing the *HIS3* locus with G418 or nourseothricin resistance markers. The *HIS3::KanMX* cassette was amplified from the strain 78 using primer pairs HIS3-F/HIS3-R. The *HIS3::NatMX* locus was amplified by a two-steps fusion PCR: the flanking regions of the *HIS3* locus were amplified from the strain LA128 using HIS3-F/HIS3-fusionTermF and HIS3-R/HIS3-fusionProm-R while the *NatMX* cassette was amplified from the plasmid pAG36 using HIS3Fusion TEFProm-F/HIS3Fusion TERterm-R. pAG36 was a donation from John McCusker (Addgene plasmid # 35126 ; http://n2t.net/addgene:35126;RRID:Addgene_35126) (Goldstein and McCusker, 1999).

The transformation of parental strains with the *NatMX* or *KanMX* cassettes was performed by electroporation as described by (Di Rienzi et al., 2011) using 1 μg of DNA and electroporation at 1.5 kV, 25 μF and 200 Ω using a GenePulser (Biorad). To confirm successful replacement of the *HIS3* locus, colonies were patched on SC-His-Ura-Thr medium and colony PCR were performed using primer pairs HIS3-F/Kan-R or HIS3-F/Nat-R. All primers used are detailed in Table S6.

### Mating, sporulation and spore isolation

For mating, LA128 *HIS3::KanMX* and LA136*HIS3::NatMX* were mixed on DYM plates for 72 h at 22°C. Double resistant cells to G418 and nourseothricin were selected on YPD-agar-G418-Nourseo plates and single colonies were purified by striking on YPD-agar-G418-Nourseo plates. The diploid state of the hybrids LA128 *HIS3::KanMX* / LA136 *HIS3::NatMX* was verified by flow cytometry. After ploidy validation, one hybrid was selected and sporulated for 2-3 days on DYM plates at 22°C. Tetrads dissections were performed using the SporePlay (Singer Instrument) without any pre-treatment. Dissection of about 1,000 tetrads showed 2:2 segregation of the KanMX and NatMX markers.

### Flow cytometry

The cell DNA content of LA128 *HIS3::KanMX* / LA136 *HIS3::NatMX* hybrids was measured by flow cytometry after synchronization with hydroxyurea as described by (Di Rienzi et al., 2011) with minor modifications. Briefly, 0.2 mL of an overnight culture was diluted in 0.8 mL of YPD and incubated for 2 h at 30°C. Cells were collected by centrifugation, resuspended in 1 mL of YPD containing 7.6 mg/mL of hydroxyurea and incubated for 2 h at 30°C. Cells were pelleted, washed in 1 mL of water and resuspended in 1 mL of ethanol 70%, pelleted again and resuspended in 1 mL of citrate buffer (sodium citrate 50 mM pH 7.4). Then cells were centrifuged and resuspended in 0.5 mL of citrate buffer containing 1 mg/mL of RNAse A. After incubation for 1 h at 50°C, 50 μl of proteinase K at 20 mg/mL were added and the cells were incubated for an additional hour at 50°C. Cells were then sonicated for 10 sec at 20% power and 0.5 mL of citrate buffer containing 2 μM of SYTOX green (Invitrogen) was added to the suspension. Cell content was analyzed with a BD Accuri C6 plus flow cytometer (BD Biosciences).

### Phenotyping

High-throughput phenotyping was realized as described in (Fournier et al., 2019) with minor modifications. Strains were pinned onto a solid YPD matrix plate to a 1,536-density format using the replicating ROTOR robot (Singer Instruments) and grown overnight at 30°C. Then the matrix plate was replicated onto phenotyping media. For each condition, all the 421 segregants are replicated three times, the hybrid384 times and the two parents 192 times. Therefore, two 1,536-density plates per media was used for phenotyping, with at least one replicate per strain on each plate. The plates were incubated for 24 h at 30°C (except for 14°C condition) and were scanned with a resolution of 600 dpi at 16-bit greyscale. Colony size was quantified using the R package Gitter (Wagih and Parts, 2014) at 0 h and 24 h. Growth of each replicate was measured by subtracting colony size at t = 0 h by colony size at t = 24 h. Within each media, Plate effect was evaluated with hybrid replicates equally distributed on the two plates and corrected. Average of the three replicates was used for final segregants phenotypic value.

### DNA extraction

Total genomic DNA of 421 segregants was extracted using the 96-well E-Z 96 Tissue DNA kit (Omega) following a modified bacterial protocol. Cells were grown overnight at 30°C with agitation at 200 rpm in 1 mL of YPD in 2-mL 96 deep square well plates, sealed with Breath-Easy gas-permeable membranes (Sigma-Aldrich). Cells were centrifuged for 5 min at 3,700 rpm and the cell wall was digested for 2 h at 37 °C in 800 μl of buffer Y1 (182.2 g of sorbitol, 200 mL of EDTA 0.5 M pH 8, 1 mL of β-mercaptoethanol, qsp 1 L of H_2_O) containing 0.5 mg of Zymolase 20T. Next, cells were pelleted, resuspended in 225 μL of TL buffer containing OB protease and incubated overnight at 56 °C. Then DNA extraction was continued according to the manufacturer’s instructions. Total genomic DNA of hybrids was extracted using a modified MasterPure Yeast DNA purification protocol (Lucigen). DNA concentration was measured using the Qubit dsDNA HS assay (ThermoFischer) and the fluorescent plate-reader TECAN Infinite Pro200 and DNA quality was evaluated using a NanoDrop 1000 Spectrophotometer (ThermoFischer) and by agarose gel analysis.

### Genotyping sequencing

DNA libraries were prepared from 5 ng of total genomic DNA using the NEBNext Ultra II FS DNA Library kit for Illumina (New England Biolabs). All volumes specified in the manufacturer’s protocol were divided by four. The adaptor-ligated DNA fragments of about 300-bp were amplified with 8 cycles of PCR using indexed primers. A combination of 48 i7 oligos (NEBNext Multiplex Oligos for Illumina, NEB) and 24 i5 oligos (Microsynth) were designed enabling multiplexing up to 1152-samples. After quality check using a Bioanalyzer 2100 (Agilent Technologies) and quantification using the Qubit dsDNA HS assay, 4 nM of each of the libraries were pooled and run on a NextSeq 500 sequencer with paired-end 75 bp reads by the EMBL’s Genomics Core Facility (Heidelberg, Germany).

### Mapping and Single Nucleotide Polymorphisms (SNPs) calling

Sequencing reads from Fastq files were mapped to the *L. waltii* reference genome (obtained from the GRYC website (http://gryc.inra.fr/index.php?page=download) using bwa mem (v0.7.17). Resulting bam files were sorted and indexed using SAMtools (v1.9). Duplicated reads were marked and sample names were assigned using Picard (v2.18.14). GATK (v3.7.0) was used to realign remaining reads. Candidate variants were then called using GATK UnifiedGenotyper.

### Segregation analysis

After variant calling, SNPs called in the LA128 and LA136 parents were first filtered (bcftools view, v1.9) to define a set of confident markers. Positions with a single alternate allele, supported by at least 10 sequencing reads in each parent and with >90% of the sequencing reads covering either the reference or alternate allele. For each strain resulting from the LA128/136 hybrid, SNPs located at aforementioned marker positions were extracted, and parental origin was assigned based on SNP correspondence between parents and spores at those positions.

In order to validate these SNPs as markers for QTL mapping, their segregation among the progeny was investigated. If most of the markers (67 %) follow the expected 2:2 mendelian segregation, a significant amount displays other patterns. Indeed, 20 % of the markers show 0:4 / 4:0 segregation illustrating loss of heterozygosity (LOH) events in the LA128/136 hybrid. Distribution of these 0:4 / 4:0 SNPs along the genome shows that at least 13 LOH events occurred encompassing 6 of the 8 chromosomes. In total LOH events represent 1.965 Mb of the 10.9 Mb total genome size. Without any segregation these LOH segments cannot be mapped in linkage analysis and therefore have been discarded for the following analysis.

Another significant amount group of SNPs encompassing all chromosomes B and D are deviating from 2:2 segregation with a more complex pattern. By looking coverage along the genome in LA128/136 parental hybrid, an aneuploidy with a supplemental copy of chromosome B and D was identified, explaining deviation from 2:2 segregation. Therefore, this aneuploidy was inherited in some of the segregants showing heterozygosity for theses chromosomes. In order to keep only euploid strains for QTL mapping analysis, segregants heterozygous for these chromosomes were discarded from the initial pool of F1 offspring. Therefore, 421 F1 offspring were used for linkage analysis. At the end, a subset of 5542 SNPs mapped in more than 94 % of the progeny and homogeneously distributed along the genome were selected to be used as genetic marker for linkage analysis.

### Linkage analysis

The QTL mapping analysis was performed with the R/ qtl package (Broman et al. 2003) by using the Haley-Knott regression model that provides a fast approximation of standard interval mapping (Haley and Knott 1992). For each phenotype, a permutation test of 1000 permutations tested the significance of the LOD score obtained, and a 1 % FDR threshold was fixed for determining the presence of QTL (Churchill and Doerge 1994). The QTL position was estimated as the marker position with the highest LOD score among all markers above the threshold in a 30 kb window.

### Data analysis

Except synteny analysis, all the statistical and graphical analyses were carried out using R software (R Core Team 2018).

### Estimation of heritability

The *lato sensu* heritability *h*^*2*^ was estimated for each phenotype according as follows:

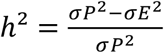

where σ*P*^2^ is the variance of progeny population in each media, explaining both the genetic and environmental variance of the phenotype measured, whereas σ*E*^2^ is the median of the variance of LA128 *HIS3::KanMX* / LA136 *HIS3::NatMX* hybrid (384 replicates per media), explaining only the environmental fraction of phenotypic variance.

### Estimation of QTL, environment and QxE effect

For each QTL mapped, the effects of QTL (Q), of environment (E) and their interactions (QxE) were estimated according to the following linear model:

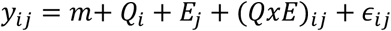

where *y*_*ij*_ is the value of the trait for allele *i* (*i* = LA128, LA136) in environment *j* (*j* = concentration level 0, …, 3), *m* is the overall mean, *Q*_*i*_ is the QTL effect, *E*_*j*_ the environment effect, (*QxE*)_*ij*_ is the interaction effect between QTL and environment and *ϵ*_*ijk*_ the residual error. The part of variance explained between *Q*_*i*_, *E*_*j*_ and (*QxE*)_*ij*_ was estimated by an analysis of variance after verification of its applicability (homoscedasticity and normal distribution of model’s residues).

### Synteny analysis

Synteny analysis was performed with the multiple genome alignment software mauve using progressiveMauve algorithm with default parameters. *L. waltii* (CBS6430, Genome Resources for Yeast Chromosomes website) and *L. kluyveri* (CBS3082, NCBI accession number: AACE00000000) genomes were used as input data.

## Data availability

All data used in this study are available in supplementary files. Genotype assignments of the segregants are available in Table S2 and their phenotypic values in Table S3.

## Figure legends

**Figure S1**. Experimental design

**Figure S2**. Distribution of QTL variance explained

## Acknowledgments

We thank members of the Schacherer lab at the University of Strasbourg for helpful suggestions throughout the project. This work was supported by the Agence Nationale de la Recherche (ANR-18-CE12-0013-02) and a European Research Council (ERC) Consolidator grant (772505). J.S. is a fellow of the University of Strasbourg Institute for Advanced Study (USIAS) and a member of the Institut Universitaire de France.

## References

Abelovska, Lenka, Marek Bujdos, Jana Kubova, Silvia Petrezselyova, Jozef Nosek, and Lubomir Tomaska. 2007. “Comparison of Element Levels in Minimal and Complex Yeast Media.” Canadian Journal of Microbiology 53 (4): 533–35. https://doi.org/10.1139/W07-012.

Ambroset, Chloé, Maud Petit, Christian Brion, Isabelle Sanchez, Pierre Delobel, Cyprien Guérin, Hélène Chiapello, et al. 2011. “Deciphering the Molecular Basis of Wine Yeast Fermentation Traits Using a Combined Genetic and Genomic Approach.” G3: Genes, Genomes, Genetics 1 (4): 263–81. https://doi.org/10.1534/g3.111.000422.

Boles, Eckhard, Patricia De Jong-Gubbels, and Jack T. Pronk. 1998. “Identification and Characterization of MAE1, the Saccharomyces Cerevisiae Structural Gene Encoding Mitochondrial Malic Enzyme.” Journal of Bacteriology 180 (11): 2875–82.

Breitling, Rainer, Yang Li, Bruno M. Tesson, Jingyuan Fu, Chunlei Wu, Tim Wiltshire, Alice Gerrits, et al. 2008. “Genetical Genomics: Spotlight on QTL Hotspots.” edited by Gary A. Churchill. PLoS Genetics 4 (10): e1000232. https://doi.org/10.1371/journal.pgen.1000232.

Brion, Christian, Claudia Caradec, David Pflieger, Anne Friedrich, and Joseph Schacherer. 2020. “Pervasive Phenotypic Impact of a Large Nonrecombining Introgressed Region in Yeast.” Molecular Biology and Evolution. https://doi.org/10.1093/molbev/msaa101.

Brion, Christian, Sylvain Legrand, Jackson Peter, Claudia Caradec, David Pflieger, Jing Hou, Anne Friedrich, Bertrand Llorente, and Joseph Schacherer. 2017. “Variation of the Meiotic Recombination Landscape and Properties over a Broad Evolutionary Distance in Yeasts.” edited by Jennifer C. Fung. PLOS Genetics 13 (8): e1006917. https://doi.org/10.1371/journal.pgen.1006917.

Broman, Karl W., Hao Wu, Śaunak Sen, and Gary A. Churchill. 2003. “R/Qtl: QTL Mapping in Experimental Crosses.” Bioinformatics 19 (7): 889–90. https://doi.org/10.1093/bioinformatics/btg112.

Churchill, G. A., and R. W. Doerge. 1994. “Empirical Threshold Values for Quantitative Trait Mapping.” Genetics 138 (3): 963–71. https://doi.org/10.1007/s11703-007-0022-y.

Clément-Ziza, Mathieu, Francesc X Marsellach, Sandra Codlin, Manos A Papadakis, Susanne Reinhardt, María Rodríguez-López, Stuart Martin, et al. 2014. “Natural Genetic Variation Impacts Expression Levels of Coding, Non-coding, and Antisense Transcripts in Fission Yeast.” Molecular Systems Biology 10 (11): 764. https://doi.org/10.15252/msb.20145123.

Dujon, Bernard. 2006. “Yeasts Illustrate the Molecular Mechanisms of Eukaryotic Genome Evolution.” Trends in Genetics. Elsevier Current Trends. https://doi.org/10.1016/j.tig.2006.05.007.

Eder, Matthias, Isabelle Sanchez, Claire Brice, Carole Camarasa, Jean Luc Legras, and Sylvie Dequin. 2018. “QTL Mapping of Volatile Compound Production in Saccharomyces Cerevisiae during Alcoholic Fermentation.” BMC Genomics 19 (1). https://doi.org/10.1186/s12864-018-4562-8.

Fabre, Emmanuelle, Héloïse Muller, Pierre Therizols, Ingrid Lafontaine, Bernard Dujon, and Cécile Fairhead. 2005. “Comparative Genomics in Hemiascomycete Yeasts: Evolution of Sex, Silencing, and Subtelomeres.” Molecular Biology and Evolution 22 (4): 856–73. https://doi.org/10.1093/molbev/msi070.

Haley, C. S., and S. A. Knott. 1992. “A Simple Regression Method for Mapping Quantitative Trait Loci in Line Crosses Using Flanking Markers.” Heredity 69 (4): 315–24. https://doi.org/10.1038/hdy.1992.131.

Janitor, Martin, and Július Šubík. 1993. “Molecular Cloning of the PEL1 Gene of Saccharomyces Cerevisiae That Is Essential for the Viability of Petite Mutants.” Current Genetics 24 (4): 307–12. https://doi.org/10.1007/BF00336781.

Kellis, Manolis, Bruce W. Birren, and Eric S. Lander. 2004. “Proof and Evolutionary Analysis of Ancient Genome Duplication in the Yeast Saccharomyces Cerevisiae.” Nature 428 (6983): 617–24. https://doi.org/10.1038/nature02424.

Kucejova, Blanka, Martin Kucej, Silvia Petrezselyova, Lenka Abelovska, and Lubomir Tomaska. 2005. “A Screen for Nigericin-Resistant Yeast Mutants Revealed Genes Controlling Mitochondrial Volume and Mitochondrial Cation Homeostasis.” Genetics 171 (2): 517–26. https://doi.org/10.1534/genetics.105.046540.

Larson, Wesley A., Garrett J. McKinney, Morten T. Limborg, Meredith V. Everett, Lisa W. Seeb, and James E. Seeb. 2016. “Identification of Multiple QTL Hotspots in Sockeye Salmon (Oncorhynchus Nerka) Using Genotyping-by-Sequencing and a Dense Linkage Map.” Journal of Heredity 107 (2): 122–33. https://doi.org/10.1093/jhered/esv099.

Lynch, Michael, and Bruce Walsh. 1998. “Genetics and Analysis of Quantitative Traits.” The American Journal of Human Genetics 68 (2): 548–49. https://doi.org/10.1086/318209.

Maharjan, Ram, and Thomas Ferenci. 2017. “The Fitness Costs and Benefits of Antibiotic Resistance in Drug-Free Microenvironments Encountered in the Human Body.” Environmental Microbiology Reports 9 (5): 635–41. https://doi.org/10.1111/1758-2229.12564.

Marullo, Philippe, Marina Bely, Isabelle Masneuf-Pomarède, Monique Pons, Michel Aigle, and Denis Dubourdieu. 2006. “Breeding Strategies for Combining Fermentative Qualities and Reducing Off-Flavor Production in a Wine Yeast Model.” FEMS Yeast Research 6 (2): 268–79. https://doi.org/10.1111/j.1567-1364.2006.00034.x.

McCouch, Susan. 2004. “Diversifying Selection in Plant Breeding.” PLoS Biology. Public Library of Science. https://doi.org/10.1371/journal.pbio.0020347.

Melnyk, Anita H., Alex Wong, and Rees Kassen. 2015. “The Fitness Costs of Antibiotic Resistance Mutations.” Evolutionary Applications 8 (3): 273–83. https://doi.org/10.1111/eva.12196.

Minikel, Eric Vallabh, Konrad J. Karczewski, Hilary C. Martin, Beryl B. Cummings, Nicola Whiffin, Daniel Rhodes, Jessica Alföldi, et al. 2020. “Evaluating Drug Targets through Human Loss-of-Function Genetic Variation.” Nature 581 (7809): 459–64. https://doi.org/10.1038/s41586-020-2267-z.

Mozhui, Khyobeni, Daniel C. Ciobanu, Thomas Schikorski, Xusheng Wang, Lu Lu, and Robert W. Williams. 2008. “Dissection of a QTL Hotspot on Mouse Distal Chromosome 1 That Modulates Neurobehavioral Phenotypes and Gene Expression.” Edited by Jonathan Flint. PLoS Genetics 4 (11): e1000260. https://doi.org/10.1371/journal.pgen.1000260.

Olson-Manning, Carrie F., Maggie R. Wagner, and Thomas Mitchell-Olds. 2012. “Adaptive Evolution: Evaluating Empirical Support for Theoretical Predictions.” Nature Reviews Genetics. Nature Publishing Group. https://doi.org/10.1038/nrg3322.

Payen, Célia, Gilles Fischer, Christian Marck, Caroline Proux, David James Sherman, Jean Yves Coppée, Mark Johnston, Bernard Dujon, and Cécile Neuvéglise. 2009. “Unusual Composition of a Yeast Chromosome Arm Is Associated with Its Delayed Replication.” Genome Research 19 (10): 1710–21. https://doi.org/10.1101/gr.090605.108.

Peter, Jackson, and Joseph Schacherer. 2016. “Population Genomics of Yeasts: Towards a Comprehensive View across a Broad Evolutionary Scale.” Yeast 33 (3): 73–81. https://doi.org/10.1002/yea.3142.

Porter, Tristan Jade, Benoit Divol, and Mathabatha Evodia Setati. 2019. “Lachancea Yeast Species: Origin, Biochemical Characteristics and Oenological Significance.” Food Research International. Elsevier Ltd. https://doi.org/10.1016/j.foodres.2019.02.003.

R Core Team. 2018. “R: A Language and Environmentfor Statistical Computing.” R Foundation for Statistical Computing, Vienna, Austria. URLhttps://Www.R-Project.Org/.

Rae, Anne M., Nathaniel Robert Street, Kathryn Megan Robinson, Nicole Harris, and Gail Taylor. 2009. “Five QTL Hotspots for Yield in Short Rotation Coppice Bioenergy Poplar: The Poplar Biomass Loci.” BMC Plant Biology 9 (1): 1–13. https://doi.org/10.1186/1471-2229-9-23.

Rienzi, Sara C. Di, Kimberly C. Lindstrom, Ragina Lancaster, Lisa Rolczynski, M. K. Raghuraman, and Bonita J. Brewer. 2011. “Genetic, Genomic, and Molecular Tools for Studying the Protoploid Yeast, L. Waltii.” Yeast 28 (2): 137–51. https://doi.org/10.1002/yea.1826.

Sharmaa, Aditi, Jun Seop Lee, Chang Gwon Dang, Pita Sudrajad, Hyeong Cheol Kim, Seong Heum Yeon, Hee Seol Kang, and Seung Hwan Lee. 2015. “Stories and Challenges of Genome Wide Association Studies in Livestock -a Review.” Asian-Australasian Journal of Animal Sciences. Asian-Australasian Association of Animal Production Societies. https://doi.org/10.5713/ajas.14.0715.

Sigwalt, Anastasie, Claudia Caradec, Christian Brion, Jing Hou, Jacky De Montigny, Paul Jung, Gilles Fischer, Bertrand Llorente, Anne Friedrich, and Joseph Schacherer. 2016. “Dissection of Quantitative Traits by Bulk Segregant Mapping in a Protoploid Yeast Species.” FEMS Yeast Research 16 (5): 56. https://doi.org/10.1093/femsyr/fow056.

Sukumaran, Sivakumar, Matthew P. Reynolds, and Carolina Sansaloni. 2018. “Genome-Wide Association Analyses Identify QTL Hotspots for Yield and Component Traits in Durum Wheat Grown under Yield Potential, Drought, and Heat Stress Environments.” Frontiers in Plant Science 9 (February): 81. https://doi.org/10.3389/fpls.2018.00081.

Supek, Frantisek, Lubica Supekova, Hannah Nelson, and Nathan Nelson. 1996. “A Yeast Manganese Transporter Related to the Macrophage Protein Involved in Conferring Resistance to Mycobacteria.” Proceedings of the National Academy of Sciences of the United States of America 93 (10): 5105–10. https://doi.org/10.1073/pnas.93.10.5105.

Vakirlis, Nikolaos, Véronique Sarilar, Guénola Drillon, Aubin Fleiss, Nicolas Agier, Jean Philippe Meyniel, Lou Blanpain, et al. 2016. “Reconstruction of Ancestral Chromosome Architecture and Gene Repertoire Reveals Principles of Genome Evolution in a Model Yeast Genus.” Genome Research 26 (7): 918–32. https://doi.org/10.1101/gr.204420.116.

Wolfgert, Hubert, Yasmine M. Manmun, and Karl Kuchler. 2004. “The Yeast Pdr15p ATP-Binding Cassette (ABC) Protein Is a General Stress Response Factor Implicated in Cellular Detoxification.” Journal of Biological Chemistry 279 (12): 11593–99. https://doi.org/10.1074/jbc.M311282200.

